# Gonad transcriptome of golden mussel *Limnoperna fortunei* reveals potential sex differentiation genes

**DOI:** 10.1101/818757

**Authors:** Luana Ferreira Afonso, Juliana Alves Americo, Giordano Bruno Soares-Souza, André Luiz Quintanilha Torres, Inês Julia Ribas Wajsenzon, Mauro de Freitas Rebelo

**Affiliations:** Biophysics Institute Carlos Chagas Filho, Federal University of Rio de Janeiro, RJ, Brazil; Bio Bureau Biotechnology, Rio de Janeiro, RJ, Brazil; SENAI Innovation Institute for Biosynthetics, National Service for Industrial Training, Center of the Chemical and Textile Industry (SENAI CETIQT), Rio de Janeiro, RJ, Brazil

## Abstract

The golden mussel *Limnoperna fortunei* is an Asian invasive bivalve that threats aquatic biodiversity and causes economic damage, especially to the hydroelectric sector in South America. Traditional control methods have been inefficient to stop the advance of the invasive mollusk, which currently is found in 40% of Brazilian hydroelectric power plants. In order to develop an effective strategy to stop golden mussel infestations, we need to better understand its reproductive and sexual mechanisms. In this study, we sequenced total RNA samples from male and female golden mussel gonads in the spawning stage. A transcriptome was assembled resulting in 200,185 contigs with 2,250 bp N50 and 99.3% completeness. Differential expression analysis identified 3,906 differentially expressed transcripts between the sexes. We searched for genes related to the sex determination/differentiation pathways in bivalves and model species and investigated their expression profiles in the transcriptome of the golden mussel gonads. From a total of 187 genes identified in the literature, 131 potential homologs were found in the *L. fortunei* transcriptome, of which 15 were overexpressed in males and four in females. To this group belong gene families relevant to sexual development in various organisms, from mammals to invertebrates, such as Dmrt (doublesex and mab3-related-transcription factor), Sox (SRY-related HMG-box) and Fox (forkhead box).

## 1. INTRODUCTION

Golden mussel *(Limnoperna fortunei)* is among the most aggressively invasive freshwater species in South America. Since its arrival from Asia in 1991, the species has spread more than 5,000 km upstream Rio de la Prata (Argentina) (1), reaching 40% of the hydroelectric power plants in Brazil and representing a loss of around USD 120 million per year (2). Golden mussel causes aquatic imbalances such as increase of cyanobacteria toxic blooms (3), grazing on phyto- and zooplankton, and the introduction of new fish parasites (4). With its high filtration rates and rapid dispersion, if prevention and control measures are not applied immediately, this mussel will likely invade most Brazilian basins, including the Amazon river (5). This current threat reflects the failing attempts to control the mussel infestation.

The success of the golden mussel as an invasive species is partially explained by its reproductive capacity. The species has at least three reproduction cycles per year and a short life span (around 3 years). The juvenile stage consists of a planktonic (veliger) larvae, which facilitates the dispersion through the water streaming (6,7). The reproductive period of *L. fortunei* is much longer than that of the most aggressive freshwater invader found in the northern hemisphere, the Zebra mussel *Dreissena polymorpha* (average of 8 months, compared to 3–4 months for the Zebra mussel) (6), with continuous production of gametes and several spawning events throughout the year (8).

Mollusks present several modes of reproduction, including strict gonochorism, simultaneous or sequential hermaphroditism, with species capable of sex change (9). Golden mussel, however, is dioecious with rare records of hermaphroditism and no external sexual dimorphism (10). With the exception of the oyster *Crassostrea gigas*, for which a working model has been suggested (11) little is known about the molecular mechanisms related to sexual development in bivalve mollusks. Most bivalve species lack heteromorphic sex chromosomes (9) and unpublished results from our research group show that this is also the case for the golden mussel. It is believed that sex determination and differentiation in this class is oligogenic and the male:female ratio may vary according to stimuli received from the environment (12).

In such species, gene expression regulation is at the core of the processes of sex determination and differentiation. Genes with sexually dimorphic expression are called sex-biased genes and can represent a high percentage of genes even in species that lack sex chromosomes (13). The identification of sex-biased genes can be achieved by comparative genome-wide gene expression analysis of male and female individuals, specially of their gonads, where there are greater functional differences between sexes. Despite the diversity of reproduction mechanisms across species, gonad transcriptome studies in bivalves suggest that many sex-determining genes are conserved across taxa and that many of these genes display sex-biased expression in the gonads, suggesting they might have a role in sex differentiation (14,11,15,16,17,18).

Previous research on sex-determining pathway genes has focused on homologous sequences of model organisms. However, identification and expression of genes related to these processes have been studied in some bivalves, including the vertebrate female-determining gene forkhead box L2 (FoxL2) (19,20,14,21), the male-determining genes double-sex- and mab-3-related transcription factor (Dmrt) (18,20,21), and Sox (Sry-related HMG box) family genes (11,12,15,16,22,23). FoxL2 and Dmrt are the key genes for sex determination and maintenance, respectively, and their antagonic role in the control of gonadal sex has been characterized in mammals and has been suggested to play similar antagonic roles in bivalve species (12,24). Some of those aforementioned genes usually present sex-biased expression in bivalve gonads, such as SoxH/Sox30 (11,12,22) and Dmrt-like (14,15,23) are male-based and FoxL2 is female-biased (11,14,16,20–23). Additionally, other conserved sex-determining related genes from model species have been identified in bivalves, such as Wnt4 (wingless-type MMTV integration site family, member 4 (11,15,16), -catenin) (11,15,16,25), and Rspo1 (R-spondin1) (15,16), among others.

Despite the identification of these several sex-determining genes, bivalve sex determination and differentiation are still poorly understood. The golden mussel genome was previously sequenced by our group (19), and transcriptome sequencing was performed (20), however sex-biased gene expression analysis was not done. In this study, we performed RNA-seq of adult golden mussel male and female gonads, identified differentially expressed genes in each sex, and prospected the genes potentially related to sex differentiation in this species. Our work contributes to a broader understanding of sex determination and differentiation in golden mussel and reveals candidate target genes for the development of biotechnological strategies to control its reproduction in order to stop *L. fortunei* infestations.

## 2. MATERIALS AND METHODS

### A. Mussels sampling, gonad processing, and histology

Golden mussels were collected from the Chavantes reservoir, Paranapanema river, São Paulo State, Brazil. *L. fortunei* is an exotic species in South America and is not classified as endangered or protected species. The gonad from each specimen was removed and divided; one part was placed in fixation buffer (4% paraformaldehyde, Phosphate Buffer 0.1 M, pH 7.4) and the other was stored in RNAlater (Invitrogen) until further pro-cessing. Fixed samples were dehydrated by increasing concentrations of ethanol, clarified with xylene, and impregnated in Paraplast Plus® (Sigma Aldrich). Histological sections of 5 μm (thickness) were submitted to Hematoxylin Eosin (HE) staining and examined under a Pannoramic MIDI slide scanner microscope (3D Histech) for gender and gonadal development stage assessment according to Callil et al. 2012 (28).

### B. RNA extraction and library preparation

RNA extraction, cDNA library construction, and sequencing were performed by a third-party company (Novogene Corporation Inc, California, US). Seven samples (four males: M2, M18, M27, and M29; and three females: M10, M15, and M17) of golden mussel gonad tissue (stored in RNAlater) at the same gonadal developmental stage (spawning) were shipped to the sequencing facility. RNA was extracted using TRIzol reagent (Invitrogen, US). Standard guidelines (29) and RNA quality control were performed by spectrophotometric analysis (NanoDrop, Thermo Fisher Scientific, US), agarose gel electrophoresis (1% concentration at 180 Volts, 16 minutes), and capillary electrophoresis using the Agilent 2100 Bioanalyzer system (Agilent Technologies, US). A total of 1 μg of RNA was used for strand-specific cDNA library construction using a NEB directional RNA lib prep kit (cat E7420L, NEB) according to the manufacturer’s protocol. The resulting 250–350 bp insert libraries were quantified using a Qubit 2.0 fluorometer (Thermo Fisher Scientific) and quantitative PCR, and size distribution of resulting fragments was analyzed with Agilent 2100 Bioanalyzer.

### C. Transcriptome sequencing, trimming, and quality control

Qualified libraries were sequenced on an Illumina HiSeq 4000 Platform with a paired-end 150 run (2×150 bases). The raw reads were submitted for quality control using the FastQC (http://www.bioinformatics.babraham.ac.uk/projects/fastqc/) of all the datasets (i.e., RNA-seq forward and reverse reads of gonad tissue of female and male golden mussel specimens). RNA-seq trimming was performed with Trimmomatic software (TruSeq3-PE.fa:2:30:10 SLIDINGWINDOW:4:5 LEADING:5 TRAILING:5 MINLEN:25). The raw data are accessible from the NCBI Short Read Archive through the BioProject accession number: PRJNA587212.

### D. Transcriptome assembly and quality assessment

We used three strategies to assemble the transcriptome of golden mussel gonads. These strategies were: *de novo* construction using Trinity (version 2.8.4) software (30), and genome-guided assembly using Trinity and StringTie (version 2.1.0) (31). De novo assembly was performed using all RNA-seq samples. For genome-guided assemblies, we used the HISAT2 v2.1.0 (32) package for mapping the reads of RNA-seq (M2 to M27) samples against the reference genome of the golden mussel, which is available under accession number GCA_003130415.1. The SAM-file produced by HISAT2 was converted into BAM and classified using SAMtools (version 1.9-45) (33). The alignments generated by the HISAT2 were then used to assemble the transcriptomes using StringTie or Trinity. To reduce redundancy while keeping the relevant biological information we used the script concatenator.pl (34) with the three separate assemblies as input. The script performed intra-assembly clustering with CD-HIT-EST (35), the unique transcripts from the assemblies were concatenated, and ORFs were predicted using TransDecoder ver.3.0.1 (http://transdecoder.github.io). These ORFs were re-clustered based on their sequence similarity, yielding high-quality final transcripts, referred to as the concatenated transcriptome assembly. We used the AssemblyStats within the Galaxy platform (36) and Trinity scripts (TrinityStats.pl) to calculate the basic assembly statistics (see Table 1 for a complete list). We also used of BUSCO (Benchmarking Universal Single-Copy Orthologs) (37) analysis and the alignment of RNA-seq reads against the assembled transcriptome using the Bowtie2 (38) software as quality metrics of the transcriptome assembly.

**Table 1.** Summary statistics of the sequencing and transcriptome assembly of gonadal tissue of *L. fortunei*

|  | *De novo* assembly<br>Trinity | Genome-guided<br>Trinity | Genome-guided<br>StringTie | Concatenated<br>assembly |
| --- | --- | --- | --- | --- |
| Lengths (bp) |  |  |  |  |
| Min contig length | 169 | 178 | 200 | 297 |
| Max contig length | 142,411 | 27,519 | 26,317 | 139,030 |
| Mean contig length | 681.73 | 677.00 | 911.11 | 1369.52 |
| Standard deviation | 1,169.99 | 792.34 | 1,108.22 | 1,988.89 |
| Median contig length | 378 | 410 | 518 | 764 |
| N50 contig length | 985 | 922 | 1,477 | 2,254 |
| Number of contigs |  |  |  |  |
| Total number of contigs | 1,673,662 | 813,347 | 332,705 | 200,185 |
| Number of contigs $\geq 1$kb | 240,281 | 126,693 | 83,050 | 79,636 |
| Number of contigs in N50 | 244,554 | 141,522 | 51,735 | 32,244 |
| Number of bases |  |  |  |  |
| Bases in all contigs | 1,140,984,094 | 550,632,535 | 303,132,232 | 274,157,883 |
| Bases in contigs $\geq 1$kb | 566,253,836 | 261,083,283 | 189,437,642 | 208,373,417 |
| GC (%) | 35.00 | 33.96 | 34.56 | 37.38 |

### E. Transcript annotation

The functional annotation took as input the nucleotide sequences of the transcripts (200,185 transcripts) generated in the final concatenated assembly using: i) similarity searches against Uniprot KB/Swiss-Prot (Release 2018_11 de 05-Dez-2018; 39) and NCBI NR (40) applying the BLASTx and Diamond algorithms (E-value < 10-3), respectively; ii) annotation with KEGG orthology and pathways through the KAAS web server (41) with the BBH (bi-directional best hit) method and the BLAST search; and iii) Gene Ontology (GO) terms assignment through Blast2GO (version 5.2) (42) using as input the resulting BLAST hits against UniprotKB/Swiss-Prot and NR NCBI databases. Conserved domains search against Pfam and CDD databases using InterProScan (version 5.33) (43) was performed with the putative protein sequences (224,899) as input.

### F. Differential transcript expression analysis

Transcript expression analyses were performed using the scripts of the Trinity software that uses the R Bioconductor repository. The transcripts from the concatenated transcriptome assembly with less than 3 TPM (Transcripts Per Kilobase Million) were filtered out using filter_low_expr_transcripts.pl script. Clean reads from three males and three females were used for quantification with Salmon-quasi (44) followed by a calculation of expression levels using the Expectation-Maximization algorithm embedded in the Trinity differential expression modules (align_and_estimate_abundance.pl). Then, the expression counts matrix generated from another Trinity script (abundance_ estimates_to_matrix.pl) was used with the EdgeR method to perform the statistical tests between male and female groups.

### G. Enrichment analysis

GO enrichment analysis was done by applying Fisher’s exact test with Blast2GO software (FDR = 0.01, GO IDs, Biological Process). The following groups were used for the statistical test: i) all transcripts blasted against UniprotKB/Swiss-Prot database and annotated with at least one GO term as the reference set, and i) sex-biased expressed transcripts (FDR < 0.001 and |logFC| > 2) for each test group (overexpressed transcripts in males and females).

### H. Protein alignment and phylogenetic analysis

Amino acid sequences were aligned with ClustalW (45) multiple alignment. Domain structures of selected proteins were determined using SMART (http://smart.embl-heidelberg.de/). The phylogenetic tree was constructed using the neighbor-joining (NJ) method and JTT substitution matrix in MEGAX (46) with bootstrap replicates of 1000. All the accession numbers of reference sequences for domains alignment and phylogenetic analyses are listed in Supplementary Table S1.

## 3. RESULTS AND DISCUSSION

### A. Transcriptome assembly quality assessment

Our study generated a total of 136.6 Gb of raw data and each sample had 62.4 ± 5.7 million sequenced reads. Table 1 displays the metrics regarding contigs count and length of the final concatenated contigs compared to the three separate assembly strategies, revealing the high quality of the assembly transcriptome obtained. The 200,185 assembled contigs had a mean length of 1,369.52 bps with N50 of 2,254 bps.

Assembled contigs completeness assessment by the representation of 978 core metazoan genes using BUSCO found 99.3% complete recovered genes for the final *L. fortunei* transcriptome, which is the highest percentage found for completeness of a transcriptome assembly in bivalves (47–53). Additionally, 87.94% of the clean RNA-seq reads were successfully mapped back to the assembled transcriptome, showing most of the reads being represented in the final assembly. It is also possible to highlight the percentage of transcripts sharing similarities against well-known databases: from a total of 200,185 contigs sequences, 109,439 (54.7%) showed hits with curated proteins of UniprotKB/Swiss-Prot database and 138,793 (69.3%) against NCBI Non-redundant (Nr) database, of which more than 64% of the hits correspond to bivalve mollusk sequences mainly from the genera *Crassostrea*, *Mytilus*, and *Mizuhopecten*. At least one Gene Ontology term and one KEGG orthology group were mapped to 26,141 (13.1%) and 19,258 (9.6%) of the transcripts, respectively. A total of 108,855 (48.4%) out of 224,899 predicted protein sequences had at least one conserved domain family from Pfam or CDD databases. Out of all the assembled transcripts, 49,856 (24.9%) were not annotated by any of the above strategies.

### B. Sex-biased expressed transcripts

Principal Component Analysis (PCA) revealed that the main variation in transcripts expression corresponds to the sexes (55.3%), which was expected and desirable for sex-biased expressed transcripts identification in this study (Supplementary Figure S1). A total of 3,906 differentially expressed transcripts were identified (FDR < 0.001 and |logFC| > 2) between male and female gonads of golden mussel specimens (Supplementary Table S2). Of these, 2,661 (68.1%) are more expressed in males whereas 1,245 (31.9%) are more expressed in females. This prevalence of male-biased transcripts was similarly observed in other bivalve mollusks, such as the yesso scallop *Patinopecten yessoensis* (22) and the Manila clam *Ruditapes philippinarum* (12), but it differs from the Pacific oyster *C. gigas* (11) and other recent study with *P. yessoensis* (54). However, Shi and collaborators (2018) (55) observed this male-biased pattern in 19 out of 21 analyzed species from several other animal taxa, including mice (*Mus musculus*), zebra fish (*Danio rerio*), *C. elegans* and *Drosophila* species. This widely observed trend is believed to be the result of a stronger sexual selection toward males, a hypothesis known as “male sex drive” (56), which leads to a “masculinization” of animal genomes.

Enriched Gene Ontology terms for each group of differentially expressed transcripts (males and females) showed that while among the female-biased transcripts we identified 82 enriched GO terms, for male-biased transcripts we detected 19 GO terms within the Biological Processes category. In females, enriched GO class of lipid metabolism and response to endogenous metabolism were 6.80% and 4.37%, respectively. The most predominant enriched GO class category in males that differed from females was the carbohydrate metabolism accounting for 10.2% of the terms. Among female-biased transcripts, GO terms related to organic compound metabolism, response to endogenous stimulus, and cell cycle processes are significantly enriched. Within male-biased enriched GO terms, carbohydrate biosynthesis and metabolic-related processes were found, as well as a nucleoside diphosphate metabolic process which can involve cell proliferation, differentiation, and development mechanisms.

Each group of statistically sex-biased expressed transcripts that was mapped to a KEGG orthology was evaluated through KEGG Mapper (https://www.genome.jp/kegg/tool/map_pathway1.html) and 274 and 293 pathways were allocated for males and females, respectively (Figure 1). The most dominant pathways in both sexes were related to signal transduction followed by the endocrine system. Signal transduction pathways within the sex-biased transcripts might be expected, as in bivalves the reproductive regulation system is mediated by endogenous hormones or neurotransmitters (57). Moreover, the main gene families involved in sex determination and differentiation are transcription factors such as Sox, Fox, and Dmrt, which also may justify the high activity of signaling cascades for the processes identified in the golden mussel.

**Fig. 1.**
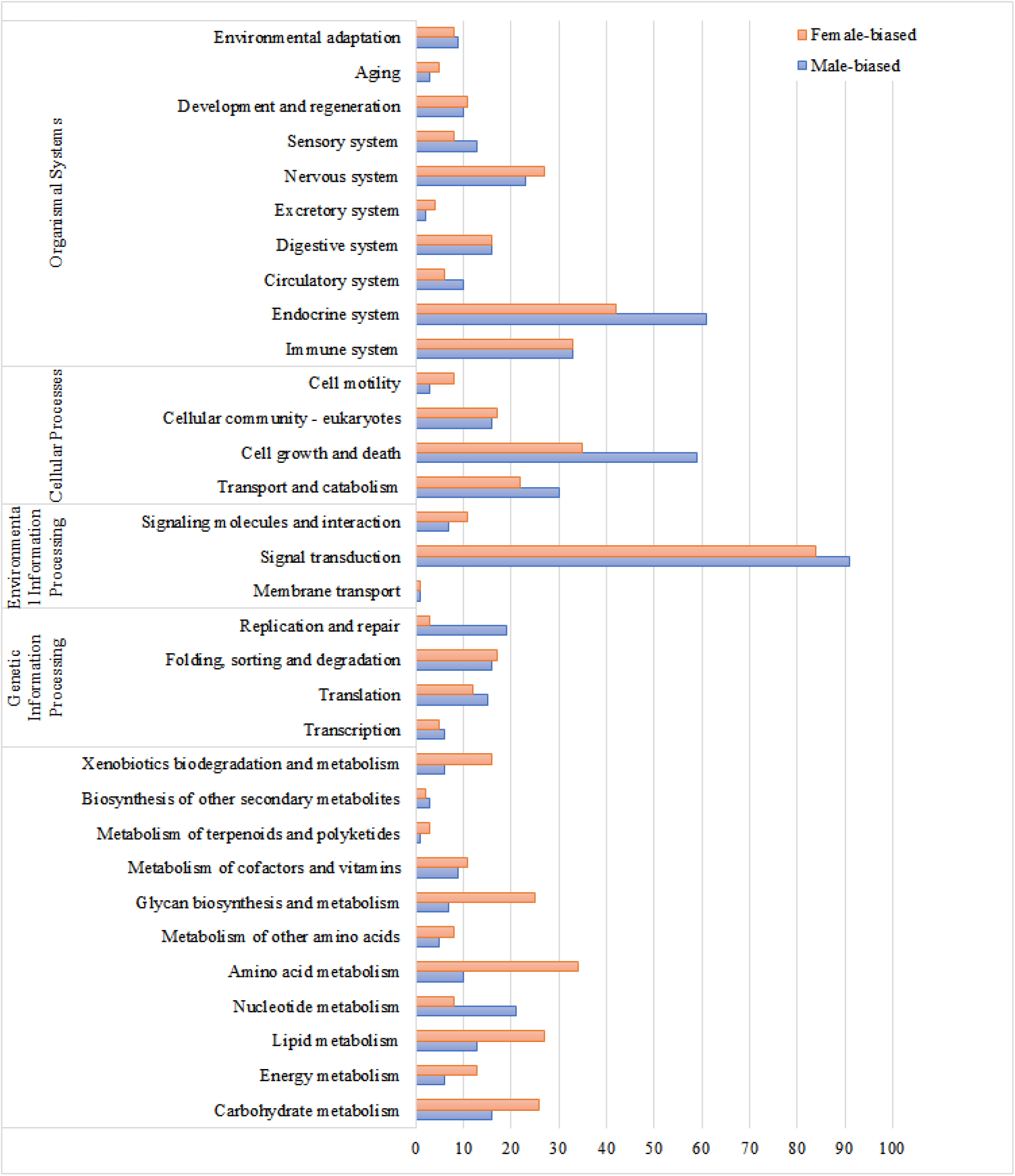
The number of differentially expressed transcripts in each sex and presented according to KEGG orthology.

### C. Identification of potential sex differentiation genes in golden mussel

To identify genes potentially involved with sex differentiation in *L. fortunei*, we used the BLASTx and Diamond results to search for genes previously associated to these processes in bivalves and model organisms (11,14–16,18,22,51,58) and investigated their expression profile in male and female gonads. Similar to Zhang and collaborators (2014), we assumed that if a gene is involved with sex-determining mechanisms in other organisms, especially bivalves, and it shows sex-biased expression according to its known function, then it may be related to these processes in *L. fortunei*. Of 187 genes related to sex determination and differentiation found in the literature, we identified 131 possible homologs in *L. fortunei* (Supplementary Table S3). Among these, the expression of 15 were male-biased whereas 4 were female-biased (Table 2).

**Table 2.** Sex-biased expression transcripts (|logFC| > 2 and FDR < 0.001) possibly involved in sex-determining processes in golden mussel. *All listed transcripts presented male-biased expression, except for Forkhead Box L2 and E4-like, Glutathione peroxidase 2 and Vitellogenin. ** Male-biased transcript with FDR < 0.05

| Transcripts ID | Description | BLAST best hit |  |  | Differential expression |  |
| --- | --- | --- | --- | --- | --- | --- |
| | | Homolog | E value | Identity (%) | $\vert \log FC\vert $ | FDR |
| GGt_738537_c0_g1_i1* | Forkhead Box L2 | *C. gigas* | 2.2E-88 | 62.05 | 8.15620 | 5.13E-15 |
| DNt_453118_c0_g1_i1* | Vitellogenin | *M. nobilis* | 0.0E0 | 52.55 | 6.45767 | 4.13E-18 |
| DNt_796766_c0_g1_i1* | Forkhead box protein E4-like | *X. laevis* | 4.1E-32 | 80.68 | 6.17922 | 1.64E-11 |
| DNt_1385095_c0_g1_i1* | Glutathione peroxidase 2 | *P. pygmaeus* | 1.9E-75 | 80.87 | 3.41515 | 6.70E-06 |
| GGt_299830_c0_g1_i1 | Male abnormal 3 | *C. elegans* | 9.0E-8 | 41.67 | 6.12615 | 1.41E-04 |
| DNt_499895_c0_g1_i1 | Meiotic recombination protein REC8 | *P. yessoensis* | 5.3E-107 | 56.76 | 4.31581 | 1.19E-05 |
| DNt_1013191_c0_g1_i1 | Meiotic recombination protein SPO11 | *C. virginica* | 2.9E-112 | 75.43 | 2.76730 | 3.49E-04 |
| GGt_312879_c0_g1_i1 | Sperm-specific H1/protamine-like protein type 1 | *O. edulis* | 6.6E-19 | 87.50 | 8.54397 | 9.29E-09 |
| GGt_149780_c0_g1_i1 | Synaptonemal complex protein 3 | *M. auratus* | 4.2E-47 | 79.76 | 5.48858 | 1.14E-13 |
| DNt_525793_c0_g1_i1 | Testis-specific serine/threonine-protein kinase 1 | *C. gigas* | 3.5E-123 | 89.05 | 9.79905 | 1.71E-41 |
| DNt_464927_c0_g1_i1 | Stabilizer of axonemal microtubules 1-like | *P. yessoensis* | 1.4E-59 | 50.32 | 9.52973 | 1.62E-54 |
| GGs_241537_c0_g1_i1 | cAMP and cAMP-inhibited cGMP 3',5'-cyclic phosphodiesterase 10A | *H. sapiens* | 0.0E0 | 62.16 | 8.65162 | 1.19E-22 |
| DNt_688365_c0_g1_i1 | E3 ubiquitin-protein ligase MARCH3-like | *C. gigas* | 4.9E-65 | 80.61 | 8.19781 | 3.71E-05 |
| GGt_42618_c0_g1_i1 | Transcription factor RFX4 | *P. yessoensis* | 4.9E-53 | 49.91 | 5.81373 | 7.39E-09 |
| DNt_604633_c0_g1_i1 | Transcription factor E2F8 | *R. norvegicus* | 2.8E-81 | 57.83 | 3.63162 | 8.79E-12 |
| DNt_844496_c0_g1_i1 | Probable ATP-dependent DNA helicase HFM1 | *P. yessoensis* | 0.0E0 | 64.99 | 3.28125 | 1.48E-07 |
| GGs_196998_c0_g1_i1 | ATP-dependent Clp protease ATP-binding subunit clpX-like | *M. musculus* | 8.4E-128 | 63.46 | 2.61628 | 1.43E-03 |
| DNt_525173_c0_g1_i1 | Transcription factor SOX15 | *D. melanogaster* | 3.4E-11 | 64.71 | 6.02617 | 1.34E-03 |
| DNt_691725_c0_g1_i1** | Transcription factor SOX30 | *M. fascicularis* | 3.8E-20 | 49.52 | 4.25569 | 4.24E-02 |

Sry-related HMG box (Sox) transcription factors belong to a gene family that shares a highly conserved high-mobility-group box (HMG) and some of their members have long been known to be related to sexual development processes, including embryogenesis and testis development (59,60). After Sry (Sex-determining region on Y) was identified in mammals, several genes encoding the HMG domain with similarity greater than 50% to that Sry domain were named as Sox genes and classified into 11 groups (from A to K) (61). In mammals, Sry is the main promoter of sexual differentiation as it suppresses ovarian-promoting genes and activates Sox9 (Sry-box 9) within the testis-determining cascade (62).

In *L. fortunei* gonad transcriptome, a male-biased (FDR < 0.05) Sox30 transcript of 6600 nt (DNt_691725_c0_g1_i1) encoding a putative 765 aa length protein was identified. Its HMG domain has 47.89% (Evalue = 2e-25) and 47.14% (Evalue = 2e-26) identities with the corresponding domains of Sry gene of *C. gigas* and *Mus musculus*, respectively. However, it shares 61.43% identity with the HMG domain of Sox30 gene of *Mus Musculus* (Evalue = 2e-27). Phylogenetic analysis shows this protein groups in a monophyletic clade (83% of bootstrap) together with Sox30 from vertebrates (Figure 2). These results are consistent with those found in *P. yessoensis* (PySOXH) and *C. gigas* (CgSOXH)(11,22). Sox30 is the only member of group H and may play key roles in gonadal development, although this gene has been characterized in only a few species so far (63).

**Fig. 2.**
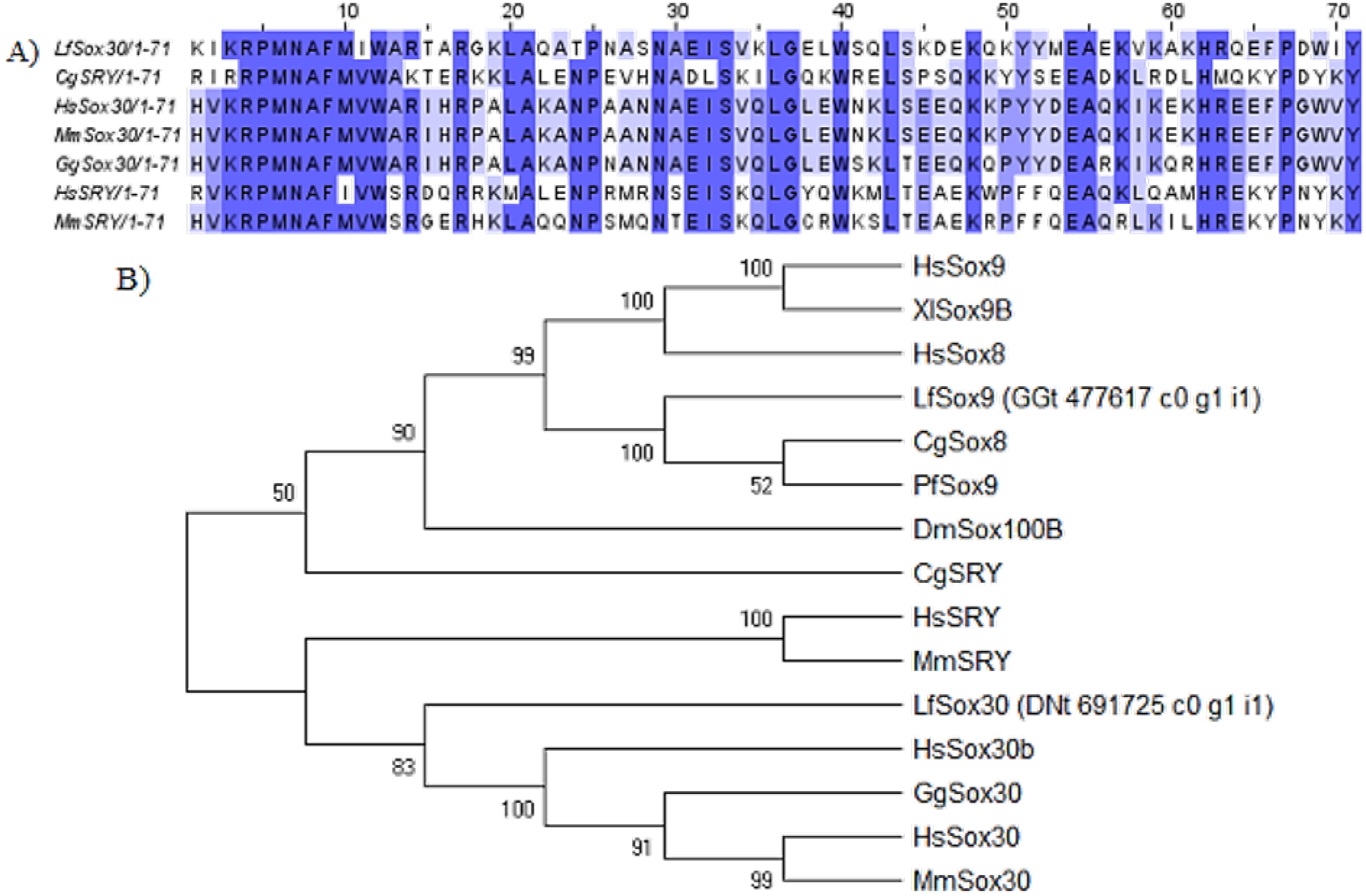
Identification of Sox genes in *L. fortunei*. (A) Alignment of HMG domains of LfSox30 and homologs from selected species. (B) Phylogenetic tree of putative protein sequences of *L. fortunei* related to Sox family genes from selected species. Species abbreviations: Hs for *Homo sapiens*, Mm for *Mus musculus*, Gg for *Gallus gallus*, Dm for *Drosophila melanogaster*, Xl for *Xenopus laevis*, Dr for *Danio rerio*, Cg for *Crassostrea gigas*, and Pf for *Pinctada fucata*.

In fact, currently only three species (*Homo Sapiens*, *Mus Musculus*, and *Macaca fascicularis*) have the Sox30 sequence curated/reviewed within the UniprotKB/SwissProt database. This gene is considered a divergent member of the Sox family and still remains under-characterized probably due to its singular classification, as its HMG domain shares low similarity with the SOX-HMG box consensus sequence (64). The Sox30/SoxH gene was found with male-biased expression in bivalves such as *C. gigas* (11), *P. yessoensis* (22,54), and *Ruditapes philippinarum* (12). The gene was also identified in the snail (*Lottia gigantea*) and the California mussel *Mytilus californianus* (data not available) (63). Another golden mussel male-biased transcript was identified as belonging to the Sox family, a Sox15-like-transcript (DNt_525173_c0_g1_i1), encoding an 882 aa putative protein. However, its HMG domain presented relatively low identity (44.29%; Evalue = 2e-20) compared to its UniprotKB/Swissprot best hit, Sox15 from *Drosophila melanogaster* (DmSox15). Phylogenetic analysis (data not shown) revealed that the protein sequence of the putative LfSox15 grouped with DmSox15 and Sry-like from *C.gigas* with low bootstrap (40-42%). Therefore, further investigation should confirm this sequence classification. In mammals, Sox15 is the only member of group G of the Sox family, being orthologous to zebrafish (*Danio rerio*) Sox19a/b and *Xenopus* SoxD, but these genes have little functionality overlap, ranging from mammalian placental cell differentiation to hair/bristle formation in different model species (65).

Another important member of the Sox family is the Sox9 gene, the only known target of Sry that participates in male differentiation and has been shown to be critical for male fertility maintenance in mammals (60). Sox9 (along with Sox8 and Sox10) belongs to the SoxE group of the Sox gene family. Sox9 was identified in *L. fortunei* as a transcript of 2029 nt (GGt_477617_c0_g1_i1), encoding a putative protein of 433 aa. The protein contains an HMG domain sharing 98.59% similarity (E-value = 1e-53) with CgSoxE (from *C. gigas*), 97.18% (E-value = 6e-53) with PfSox9 (from *Pinctada fucata*), and 94.37% (E-value = 7e-52) with HsSox9 (from *Homo sapiens*) domains, revealing that the LfSox9 HMG domain is highly conserved (Supplementary Figure S2). According to phylogenetic analysis, the *L. fortunei* Sox9 grouped with other proteins from the group E, showing closer relation to the bivalve sequences (Sox8 and Sox9 from *C. gigas* and *P. fucata*, respectively) with high bootstrap support (100). Consistent with other studies done with mollusks (11,23,25), Lf-Sox9 did not show male-biased expression, as the expression of this transcript was similar in males and females, i.e., unbiased expressed. This suggests that Sox9 gene may not be related to sex differentiation mechanisms in the golden mussel. In any case, further studies should investigate the expression pattern of the Sox9 gene in earlier developmental stages. In this study, we found that the golden mussel transcriptome has transcripts similar to other Sox family genes, such as Sox17, SoxB2, Sox4, Sox5, and Sox2, which was already suggested to play key roles in spermatogenesis and testis development in adults specimens of the scallop *Chamys farreri* (66).

Fox (forkhead-box) proteins are a family of transcription factors with a DNA-binding forkhead domain that are involved in diverse biological processes including development and sexual differentiation mechanisms. FoxL2 is a member of the Fox gene family presenting a key role in ovarian determination in vertebrates. In mammals, FoxL2 is expressed in the ovary, promoting its development, while suppressing the Sox9 gene and other testis-specific genes from early embryonic gonad throughout adult individuals (67,68). In this study, a female-biased gene FoxL2 was identified as a transcript of 1594 nt long (GGt_738537_c0_g1_i1) which encodes a putative 335 aa protein with a conserved forkhead box domain. Homology analysis on the amino acid sequences of the forkhead domain shows high sequence identity with many species including the Pacific oyster C. gigas (89.01%, E-value = 1e-63) (Figure 3). FoxL2 gene has been found in several other bivalves such as *C. farreri*, *Pinctada margaritifera*,*H. shlegelii*,*Crassostrea hongkongensis*,*Tegillarca granosa*, and *Chlamys nobilis*, all showing a sexually dimorphic pattern of expression towards females (14–16,20,21,23). However, in *C. gigas* gonad transcriptome, FoxL2 was expressed in both sexes, it was highly but not specifically expressed in the ovary, with increased expression earlier during sexual development in females. This suggests an important role for CgFoxL2 in embryogenesis, but seems to be unrelated to sex determination in *C. gigas*. (11,19).

**Figure 3.**
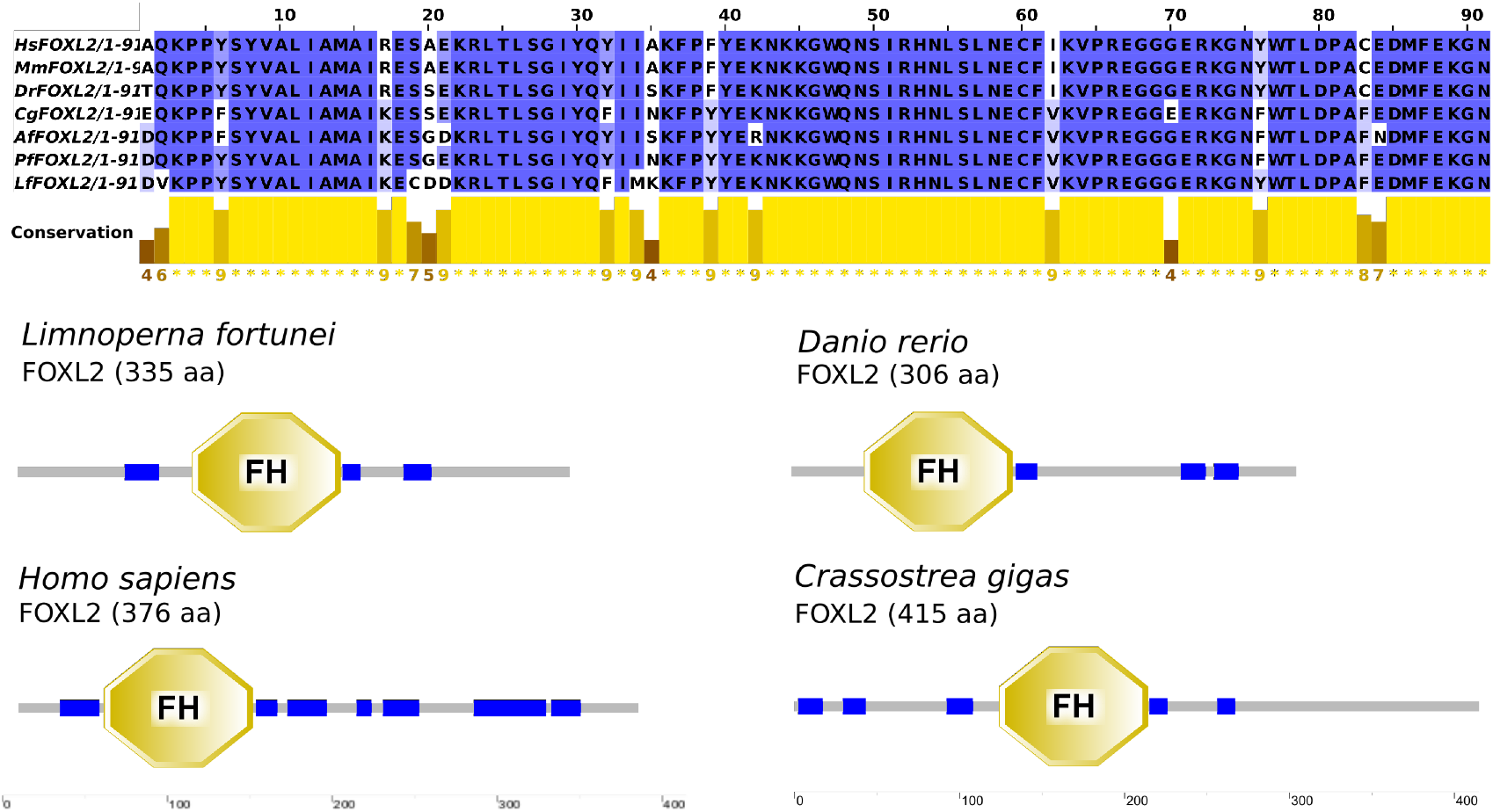
FoxL2 identification in *L. fortunei*. (A) Forkhead domain alignment of FoxL2 from *L. fortunei* and other selected species. Species abbreviations: Hs for *Homo sapiens*, Mm for *Mus musculus*, Dr for *Danio rerio*, Cg for *Crassostrea gigas*, Af for *Azumapecten farreri*, and Pf for *Pinctada fucata*. (B) FoxL2 domain structure of *L. fortunei*, *Homo sapiens*, *Danio rerio*, and *Crassostrea gigas*. Low-complexity regions are represented in blue.

The Dmrt (doublesex and mab3-related-transcription factor) gene family encodes a large family of transcription factors characterized by the presence of a DM domain DNA binding motif, and has the most conserved function among different phyla (69). This family play roles in sex determination, sexual dimorphism, and other developmental and sexual reproduction processes mostly investigated in model organisms (70). Members of the family include the doublesex (Dsx) gene in *Drosophila melanogaster*, MAB-3 in *Caenorhabditis elegans*, and the Dmrt1 in vertebrates, all participating in the male-specific development. Dmrt-like genes have been identified with male-biased expression in mollusks such as *C. gigas* (11), the abalone *Haliotis asinina* (70,71), the pearl oyster *Pinctada martensii* (72), the blacklip pearl oyster *P. margaritifera* (14), the freshwater mussel *Hyriopsis schlegelii*, and the scallops *Nodipecten subnodosus* and *P. yessoensis* (18,54). In the golden mussel transcriptome, nine putative proteins containing DM domain were found; one of them shows male-biased expression (GGt_299830_c0_g1_i1). This sequence corresponds to a 509 nt transcript encoding a putative 169 aa protein containing two DM domains. The DM domains of *fortunei* transcript share 42.55% (E-value = 4e-12) and 40.54% (E-value = 2e-08) identity with the corresponding DM domains of its best hit against UniprotKB/Swiss-Prot, MAB-3 of *C. elegans* (CeMAB-3)(Figure 4). This finding differs from that of the CgDsx (GenBank accession No. KJ489413) structure, which contains only one DM domain, however sharing 45% identity (E-value = 9e-13) with *D. melanogaster* Dsx protein (11). Another golden mussel transcript (DNt_1078970_c0_g1_i1, 3358 nt) shares potential homology with the MAB-3 of *C. elegans*, encoding a putative 316 aa protein also containing two DM domains in its structure, however this transcript showed low and unbiased expression in both male and female specimens. Although Dmrt genes are involved with a conserved role in gonad development in many animals, from vertebrates to arthropods and mollusks, DM domains show little sequence conservation, which can also obscures phylogenetic relationships (73).

**Figure 4.**
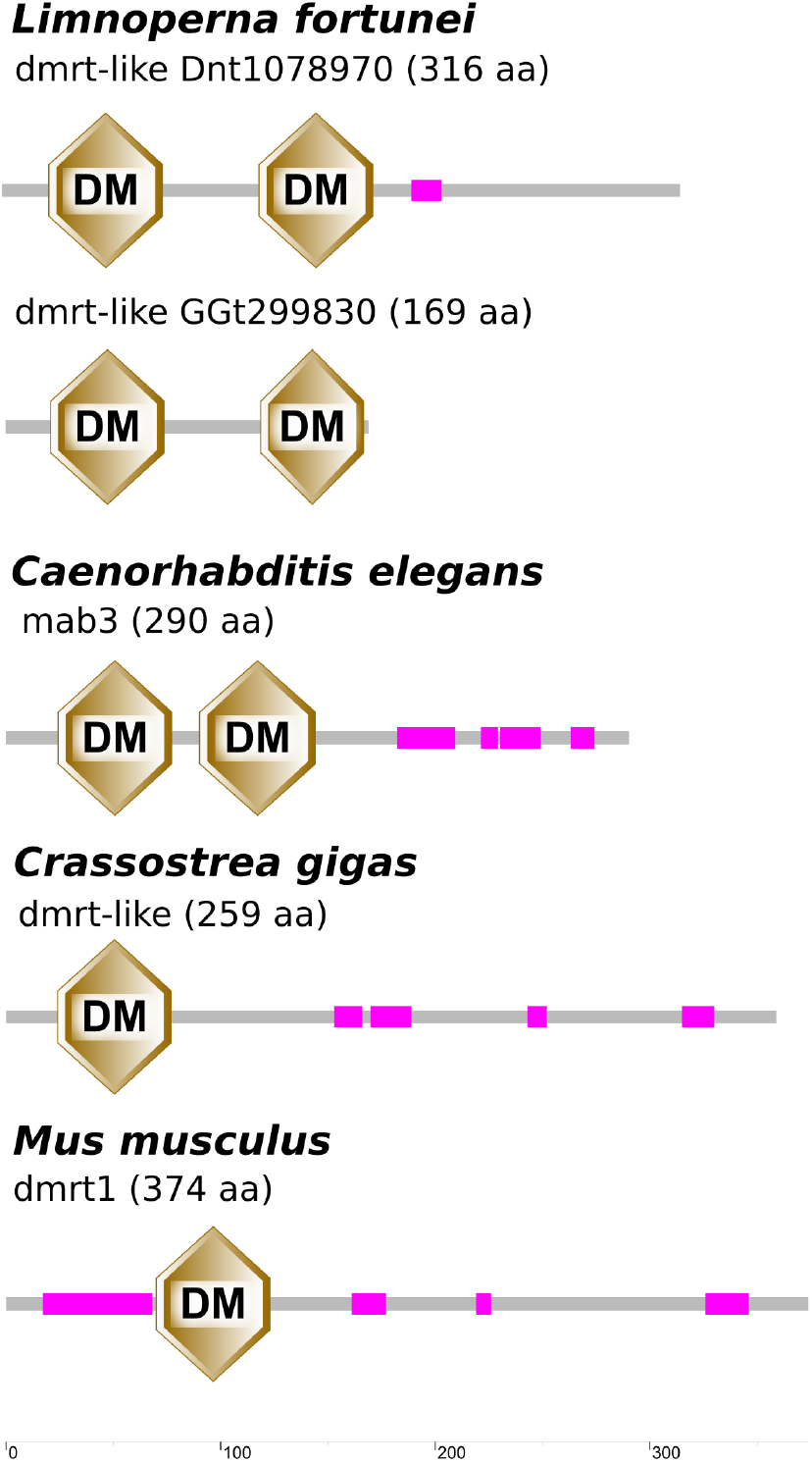
Domain structures of Dmrt-like protein of *L. fortunei* and selected DM domain genes from other species (low-complexity regions are represented in purple).

Two other Dmrt-like transcripts (GGt_460196_c0_g1_i1 and GGs_22755_c0_g1_i1) of 1012 and 1461 nt, respectively, were identified. They encode putative proteins of 295 and 351 aa and both present one DM domain and an additional DMRTA motif, a region found in the C-terminus of the DM domain. According to phylogenetic analysis, the amino acid sequence coded by GGs_22755_c0_g1_i1 grouped with DmrtA2/Dmrt5 and Dmrt4 of *C. gigas* and *P. fucata* species with high bootstrap (100%) and GGt_460196_c0_g1_i1 with Dmrt2 from mammals (*Homo sapiens* and *Mus musculus*) and invertebrates (*Danio rerio*) with bootstrap of 99% (Figure 5). Both of those transcripts presented unbiased expression patterns in both sexes in golden mussel.

**Fig 5.**
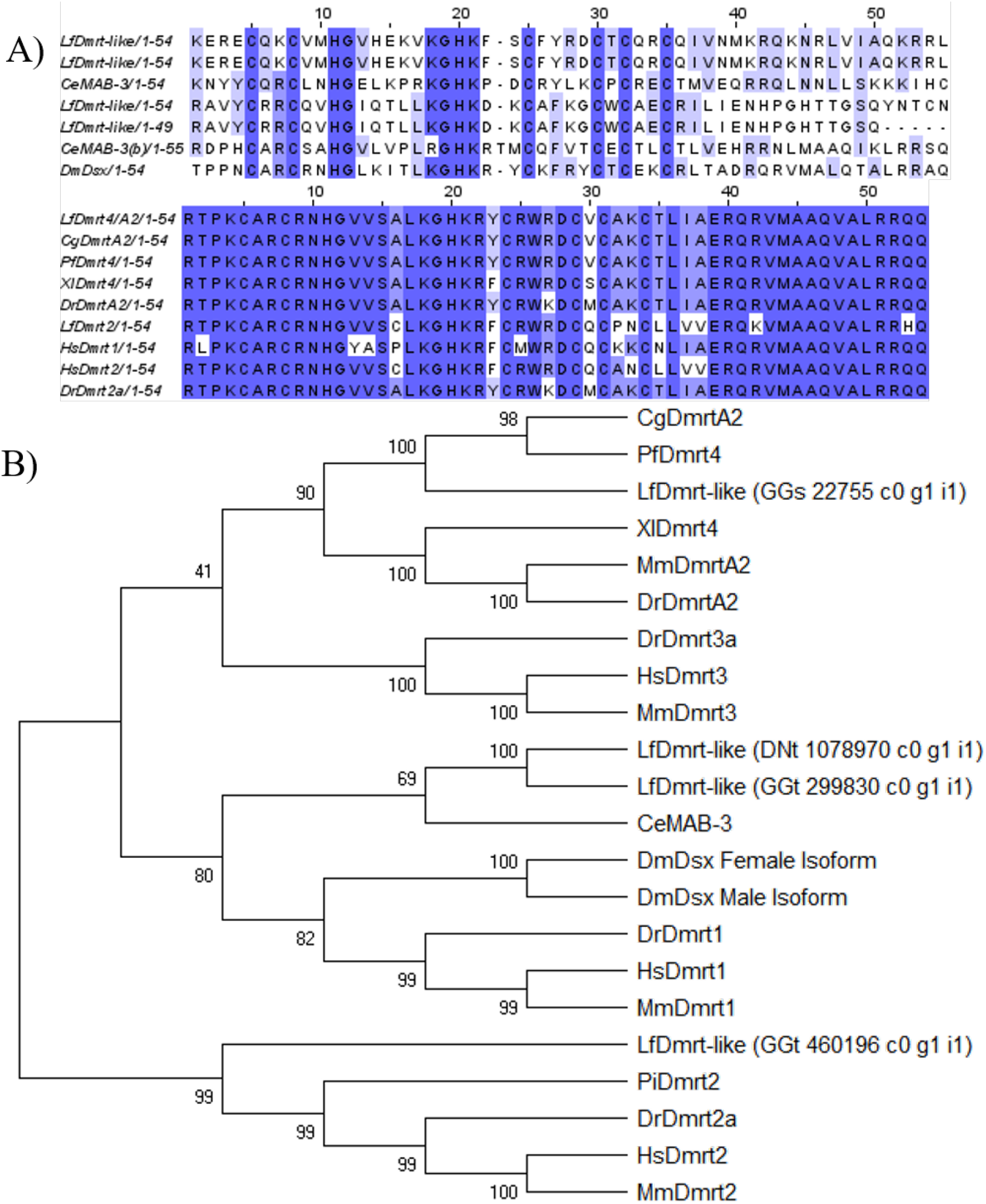
Identification of Dmrt-like sequences in *L. fortunei*. (A) DM domain alignments of *L. fortunei* amino acid sequences (LfDmrt3a: GGt_460196_c0_g1_i1; LfDmrtA2: GGs_22755_c0_g1_i1; LfDmrt-like (1): DNt_1078970_c0_g1_i1; LfDmrt-like(2): GGt_299830_c0_g1_i1) with selected species. (B) Phylogenetic tree of DM family protein sequences of *L. fortunei* with other selected species. Species abbreviations: Cg for *Crassostrea gigas*, Pf for *Pinctada fucata*, Xl for *Xenopus laevis*, Mm for *Mus musculus*, Dr for *Danio rerio*, Hs for *Homo sapiens*, Ce for *Caenorhabditis elegans*, and Dm for *Drosophila melanogaster*.

Vitellogenin was also identified with female-biased expression in golden mussel gonad transcriptome. In fact, this gene was identified in bivalve gonad transcriptome studies as being potentially involved with female development. This was observed in the pearl oyster *P. margaritifera* (14) and also in *P. yessoensis*, where it probably participates in sex determination and differentiation pathway, while Dmrt1 is repressed by ovary genes (54). Other relevant sex-determining genes described in bivalves, such as -catenin, Wnt4 (wingless-type MMTV integration site family, member 4), GATA4 (GATA binding protein 4), and Follistatin (Fst) were identified but with no significant difference in expression between males and females. However, it has been observed in other models that some genes may show a sex-biased expression pattern that depend on the developmental stage of the organism (74). Therefore, it is possible that some other genes not reported here may show a sex-biased expression pattern in developmental stages not investigated in this study. Our group is the first to publish the gonad transcriptome of the freshwater bivalve *L. fortunei*, an organism that has a highly negative impact on aquatic systems by changing the ecological environment and by colonizing and clogging pipes, causing huge economic damage.

Golden mussel seems to share key genes related to the sexual development of model organisms from vertebrates to insects and nematodes and that have been identified in other bivalve species such as *C. gigas*, *P. margaritifera* and *P. yessoensis* (11,75,14,22). Our next step is to analyze the expression levels of these genes across all developmental stages and experimentally validate their role in sex determination/differentiation processes during *L. fortunei* development.

## 4. CONCLUSION

The sequencing of the transcriptome of *L. fortunei* allowed us to identify several genes related to sexual development mechanisms already known in other species. Our results reveal that the golden mussel shares sex differentiation genes with other bivalve species and that some of the putative transcripts present sex-biased expression. These include the Sox family genes such as Sox30 and Dmrt-like showing male-biased expression, and the female-biased FoxL2. These genes are known to be associated with sexual development not only in bivalve species but in virtually all animals.

## Supporting information

Table S1. Species and accession numbers of reference sequences

Table S2. Differentially expressed transcripts from males and females gonads of Limnoperna fortunei

Table S3. Potential sex differentiation transcripts in Limnoperna fortunei

**ABBREVIATIONS**
FDR: False Discovery Rate
GO: Gene ontology
KEGG: Kyoto Encyclopedia of Genes and Genomes
PCA: Principal Component Analysis
ORF: Open Reading Frame

---

**Figure S1.**
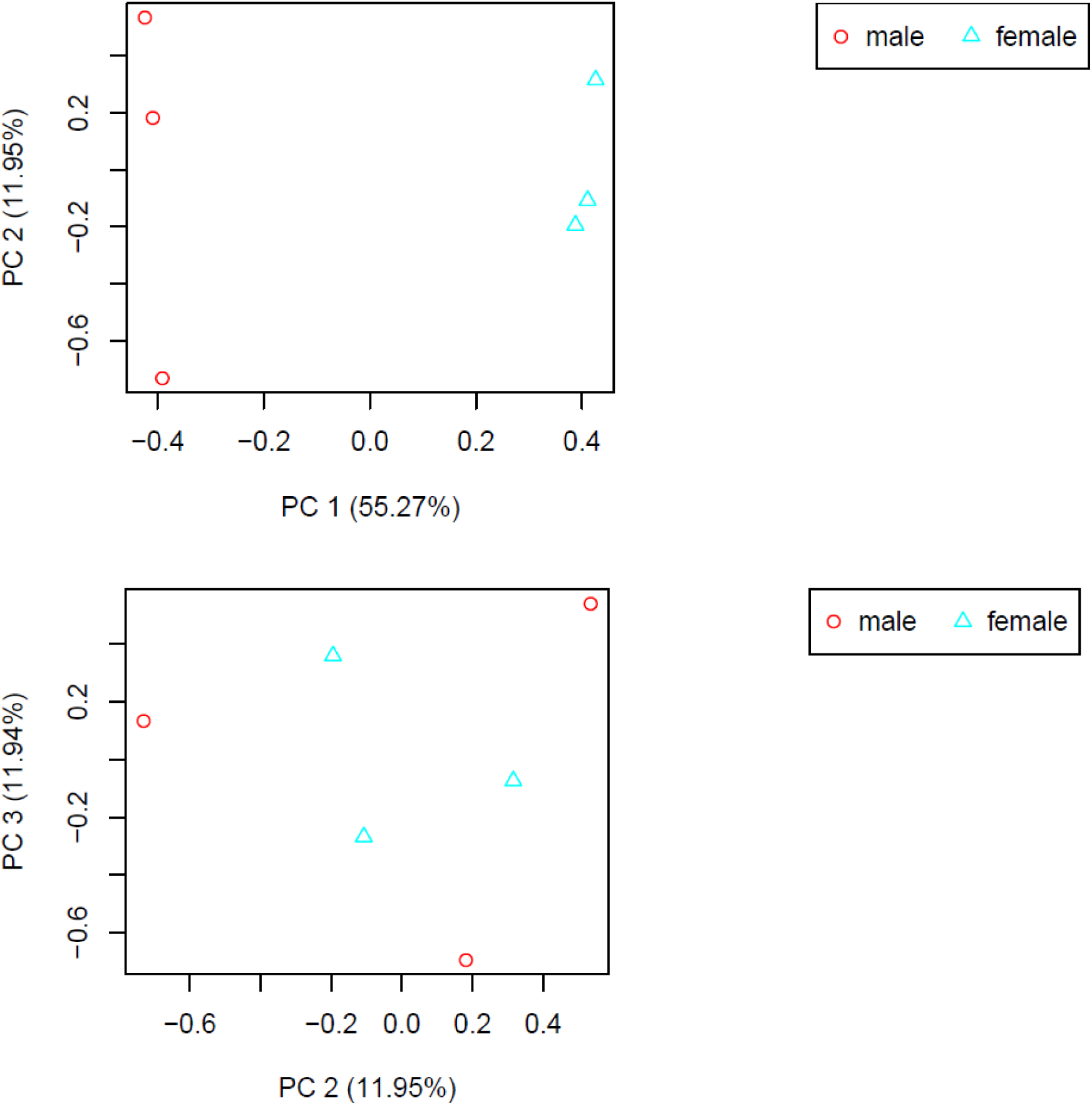
Principal component analysis (PCA) of *Limnoperna fortunei* gonads RNA-seq samples from males (red circles) and females (blue triangles) specimens.

**Figure S2.**
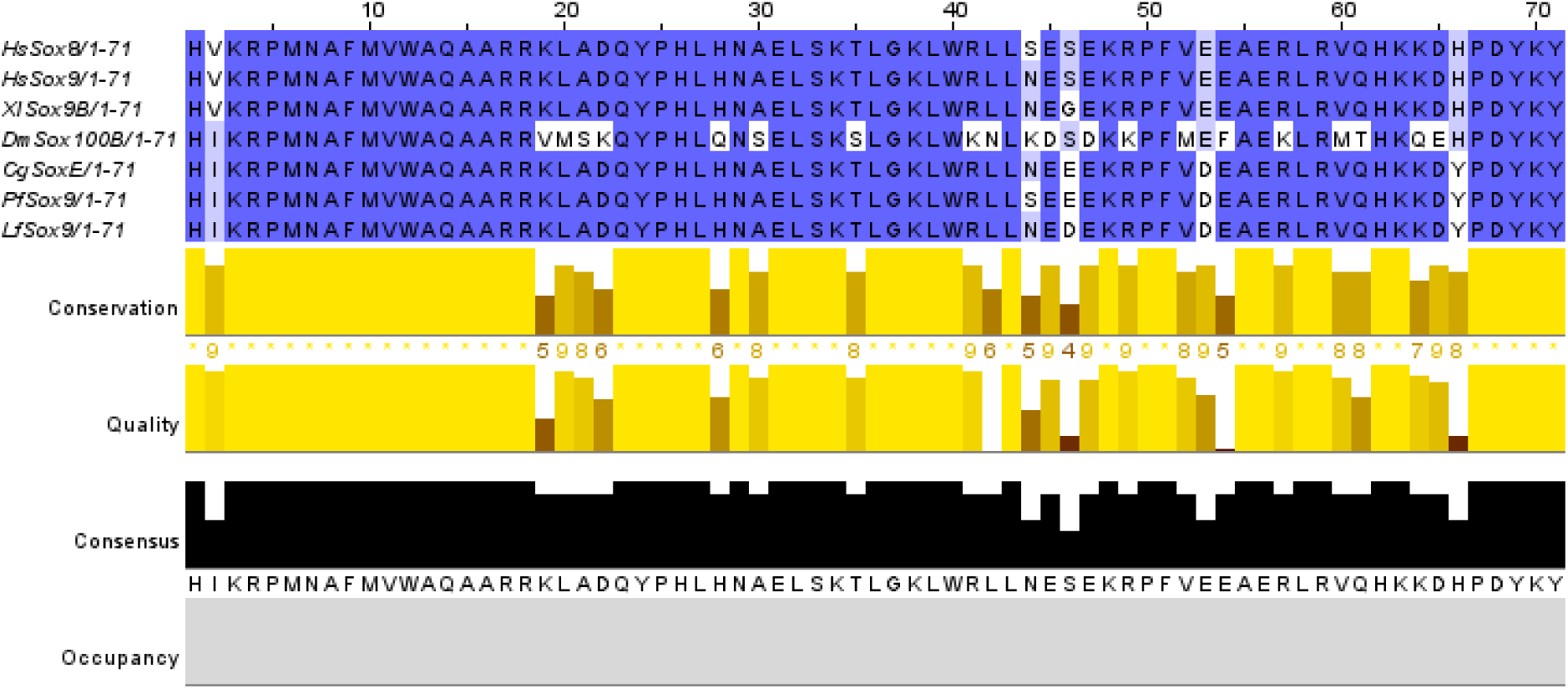
Alignment of HMG domains of *LfSox*9 and potential homologs from selected species (*Homo sapiens, Xenopus laevis, Drosophila melanogaster, Crassostrea gigas, and Pinctada fucata*)

